# A Migratory Divide in the Painted Bunting (*Passerina ciris*)

**DOI:** 10.1101/132910

**Authors:** C.J. Battey, Ethan B. Linck, Kevin L. Epperly, Cooper French, David L. Slager, Paul W. Sykes, John Klicka

**Affiliations:** University of Washington Department of Biology and Burke Museum of Natural History & Culture, Seattle, WA 98115

## Abstract

Divergence in migratory behavior is a potential mechanism of lineage divergence in sympatric populations and a key life history trait used in the identification of demographically independent units for conservation purposes. In the Painted Bunting (*Passerina ciris*), a North American songbird, populations on the Atlantic coast and interior southern United States are known to be allopatric during the breeding season, but efforts to map connectivity with wintering ranges in Mexico, Florida, and the Caribbean have been largely inconclusive. Using genomic and morphological data from natural history specimens and banded birds, we found evidence of three genetically differentiated populations with distinct wintering ranges and molt-migration phenologies. In addition to confirming that the Atlantic coast population remains allopatric throughout the annual cycle, we identified an unexpected migratory divide within the interior breeding range. Populations breeding in the Lower Mississippi River Valley winter on the Yucatán Peninsula, and are parapatric with other interior populations that winter in mainland Mexico and Central America. Across the interior breeding range, genetic ancestry is also associated with variation in wing length; suggesting that selective pressures may be promoting morphological divergence in populations with different migration strategies.

## Introduction

Migratory divides occur in regions where sympatric or parapatric populations differ in the timing or route of seasonal migration. Because migratory behaviors have clear fitness impacts and are strongly heritable in several taxa (Helbig, 1991; Pulido et al., 2001; Quinn et al., 2000; Zhan et al., 2014), migratory divides are thought to represent a mechanism of lineage divergence in sympatry (Bearhop et al., 2005; Rolshausen et al., 2009). If hybrids between populations differing in migratory behavior attempt an intermediate strategy with lower fitness than either parental type, selection is expected to favor the evolution of mechanisms that reduce the probability of breeding across migratory types (Rohwer and Irwin, 2011). Recent studies combining genetic and individual tracking data have documented this scenario in a passerine bird (Delmore and Irwin, 2014), though reduced hybrid fitness has yet to be rigorously tested in the wild.

Understanding the role of seasonal migration in mediating gene flow among populations is also important in identifying distinct evolutionary and demographic units relevant for making conservation decisions. Because the level of immigration required to homogenize allele frequencies among populations is much lower than that expected to drive trends in population size (Waples and Gaggiotti, 2006), evidence of genetic differentiation is a conservative proxy for demographic independence. Migratory connectivity has long been recognized as a core criteria for delimiting fish stocks (Cadrin et al., 2013; Gillanders, 2002; Lipcius et al., 2008), but it has not been widely used in monitoring songbird populations. In part, this flows from our relatively sparse knowledge of variation in migratory behavior within most species of bird (Faaborg et al., 2010).

The Painted Bunting (*Passerina ciris*) is a seasonal migrant to the southern United States, with an interior population that stretches from the Texas Panhandle east through Louisiana (*P. c. pallidior*), and a second population that hugs the Atlantic Coastline from southern Georgia to Virginia (*P. c. ciris*). The species winters across Mexico, Central America, Florida, and the northern Caribbean, but connectivity between specific areas of the breeding and wintering ranges remains poorly characterized. It has been suggested that the Atlantic coast population winters exclusively in southern Florida and on islands in the northern Caribbean (Storer, 1951; Thompson, 1991), although others (e.g. Sykes Jr et al., 2007) maintain that eastern birds also winter in the Yucatán and further southward. Winter destinations of interior migrants are similarly unresolved. Storer (1951) and later Thompson (1991) proposed that the birds breeding in Louisiana and Mississippi are trans-gulf migrants that winter on the Yucatán Peninsula, while birds that breed further west use a circum-gulf route to sites elsewhere in Mexico and Central America. Rohwer et al. (2013) and Contina (2013) demonstrated that Sinaloa serves as a stopover site where migrants from the interior group molt before working their way down the Mexican Pacific coast. In a metanalysis of specimen collection records, Linck et al. (2016) extended this hypothesis to suggest that these buntings migrate in a circular pattern around Mexico, tracking resource availability. A phylogeographic study across the species’ breeding range showed significant population structure between coastal and interior populations (Herr et al. 2011); however, genetic data have not yet been used to identify links between breeding and wintering grounds.

Here we use genome-wide DNA sequence data and morphological analyses of museum specimens to infer phylogeographic history and patterns of migratory connectivity in the species. Specifically, we 1) map migratory connectivity between breeding and wintering grounds, 2) test for morphological variation associated with divergent migratory strategies in the interior population, and 3) estimate divergence times and rates of gene flow among current populations. Our results highlight the contrasting roles of seasonal migration in driving both gene flow and genetic differentiation, and have significant conservation implications for an iconic but declining songbird.

## Methods

### Genetic Sampling

We collected a total of 260 blood, tissue, and feather samples from across the breeding and wintering ranges of *P. ciris* (Supplementary Table 1-2), including 138 breeding-range samples previously analyzed in Herr et al. 2011. All Atlantic Coast samples were blood and feather samples taken during banding studies. Interior populations were represented by 148 frozen tissue samples of vouchered museum specimens held by the Burke Museum of Natural History and Universidad Nacional Autónoma de México.

Whole genomic DNA was extracted using Qiagen DNEasy extraction kits. We sequenced the mitochondrial gene NADH dehydrogenase subunit 2 (ND2) from all samples using the primers and protocol described in Herr et al. 2011. Based on fragment size and DNA concentration, 95 samples were selected for reduced-representation library sequencing via the ddRADseq protocol (Peterson et al., 2012). We used the digestion enzymes sbf1 and msp1 and a size-selection window of 415-515 bp. The resulting libraries were sequenced for 100 bp single-end reads on an Illumina Hiseq 2500.

Reads were assembled into sequence alignments using *de-novo* assembly in the program pyRAD v.3-0-65 (Eaton, 2014). We set a similarity threshold of 0.88 for clustering reads within and between individuals, a minimum coverage depth of 5, and a maximum of 8 low-quality reads per site. To exclude paralogs from the final dataset, we filtered out loci with more than 5 heterozygous sites and those sharing a heterozygous site across more than 60 samples. For clustering analyses, we required each locus to be sequenced in at least half of samples and randomly selected one parsimony-informative SNP per locus using a custom R script (<u>https://github.com/slager/structure_parsimony_informative</u>).

### Genotype Clustering & Population Assignments

We used multivariate ordination and Bayesian coalescent clustering to identify genetically differentiated populations and assign wintering individuals to breeding regions. For multivariate analyses, we first conducted a principal components analysis (PCA; Pearson, 1901) on allele frequencies in each sample and then identified putative genetic clusters using k-means clustering in the R package Adegenet v2.0.1 (Jombart, 2008). We then cross-validated population assignments with discriminant analysis of principal components (DAPC; Jombart et al., 2010) by randomly selecting half the samples in each *k-*means cluster, conducting a DAPC on these samples, and predicting the group assignments of remaining individuals using the training DAPC. Cross-validation analyses were repeated 1,000 times for *k* = 2-4 to estimate cluster assignment accuracy.

Bayesian clustering under a coalescent model with admixture was implemented in the program Structure (Pritchard et al., 2000) using default priors, correlated allele frequencies, and a chain length of 1,000,000. The first 100,000 steps were discarded as burnin. Structure analyses were replicated 5 times for each value of *k=*2-4. We assessed change in marginal likelihood across values of *k* using Structure Harvester (Earl, 2012), and used CLUMPP (Jakobsson and Rosenberg, 2007) to take the mean of permuted matrices across replicates after accounting for label switching. We developed a custom web app for visualizing Structure results (<u>https://cjbattey.shinyapps.io/structurePlotter/</u>; Battey, 2017), and summarized output of multivariate analyses using the R packages ‘plyr’ (Wickham, 2015) and ‘ggplot2’ (Wickham, 2016).

### Mitochondrial DNA

Mitochondrial DNA analyses focused on identifying the most likely breeding population of wintering samples collected in areas not covered by ddRAD sequencing. We created 12 hypothetical population assignment schemes based on the results of nuclear SNP clustering, varying only the population assignment of samples from regions without nuclear SNP sequence data (Cuba, Bahamas, Costa Rica, and Nicaragua; Supplementary Table 2). Assignment schemes were compared by conducting an analysis of molecular variance (AMOVA; Excoffier et al., 1992) in the R package ‘poppr’ (Kamvar et al., 2014) and ranking models by the percentage of total variance explained by the population factor (following Herr et al. 2011). We also inferred a median-joining haplotype network using the R package ‘pegas’ (Paradis, 2010) to visualize the distribution of haplotypes among putative populations.

### Demographic Modeling

To estimate the timing of population splits and rates of gene flow among populations, we fit demographic models to the joint site frequency spectrum (SFS) of our nuclear SNP alignment in the program δaδi v1.7 (Gutenkunst et al., 2010). We randomly selected 10 samples from each *k-*means population and called SNPs from this subset in pyRAD. PyRAD VCF files were then converted to δaδi’s input format using a custom R script (see data supplement). A single biallelic SNP was randomly selected from each locus, and the final dataset was projected to a size of 5 individuals per population (proj=[10,10,10]), yielding 3,044 SNPs from 4,128 loci.

We fit two 9-parameter demographic models meant to capture the general phylogeographic history of the group (Supplementary Figure 1). In both models a single ancestral population splits into Eastern and Western groups, one of which then splits to form the Central population. Migration is allowed between east+central and west+central populations after the final divergence event. The models differ only in whether eastern or western birds are sister to the central population. We ran 40 optimizations from randomized starting positions for each model, using the “optimize_log()” function in δaδi, and assessed uncertainty across 100 parametric bootstrap replicates of our original data (sampling each locus with replacement). We ranked models by calculating the difference in Akaike Information Criterion (AIC; Akaike, 1974) for the highest-likelihood parameter set for each model.

To convert model parameters to demographic values, we used the average genome-wide mutation rate of *Geospiza fortis* (3.44x10^-9^ substitutions/site/year; Nadachowska-Brzyska et al., 2015) and a *Passerina* generation time of 1.63 years (Weir and Schluter, 2008). We estimated the effective sequence length for SNP calling by multiplying the total base pairs in our pyRAD alignment by the fraction of all SNPs incorporated in the SFS after projection.

### Morphology

Two previous studies have documented significant range-wide variation in Painted Bunting wing length; both concentrating primarily on differences between the allopatric coastal and interior breeding populations (Storer, 1951; Thompson, 1991). Here we focused on testing for morphological variation associated with the putative migratory divide within the interior breeding population, because variation in migration distance has previously been associated with wing length in both birds and butterflies (Altizer and Davis, 2010; Voelker, 2001). We measured wing chord and tarsus length (as a proxy for body size) of 59 museum specimens of adult male Painted Buntings collected in the breeding season (April-June). Wing chord was measured to the nearest 0.5mm with a metal stop ruler. Tarsus length was measured to the nearest 0.01mm with a set of digital calipers.

We calculated mean values for both traits at each unique specimen locality and plotted these on a map to visualize trends across geographic space. After initial analyses found that wing chord and tarsus length were not significantly correlated (*p*=0.32, *R*^2^=0.02), we treated these variables as independent. Following Slager et al. (2015), we used a principal components analysis implemented in R to create a synthetic variable combining wing chord and tarsus size. We conducted linear regressions to test for significant correlations between specimen longitude and each of wing length, tarsus length, and the first PC axis of wing and tarsus length. Finally, for the 21 specimens with both genomic and morphological data, we used linear regression to test for significant correlations between morphological traits and the proportion of “central” ancestry inferred by Structure.

## Results

### Sequence Assembly

Illumina sequencing returned an average of 721,942 quality-filtered reads per sample. Clustering within individuals identified 36,497 putative loci per sample, with an average depth of coverage of 17. After clustering across individuals and applying paralog and depth-of-coverage filters, we retained an average of 9,010 loci per sample. As in most studies using RADseq-style reduced-representation libraries, we observed a large effect of missing data filtering on the size of our alignments; ranging from 238 to 25,434 unlinked SNPs for an all-samples-present vs. three-samples-present cutoff (DaCosta and Sorenson, 2016; Eaton et al., 2015; Leaché et al., 2015). The final alignment used for clustering analyses included 3,615 unlinked parsimony-informative SNPs from 5,950 loci sequenced in at least 48 out of 95 of samples.

### Genotype Clustering

In *k-*means clustering, BIC showed a single clear inflection point at a *k* value of 2; while Structure returned the highest marginal likelihood at *k*=3 (Supplementary Figure 2). Individual population assignments were similar in both methods, with *k=*2 models splitting breeding individuals between the Atlantic Coast and Interior breeding ranges, and wintering individuals between Florida and Mexico + Central America (Figure 1). At *k=*3, both methods cluster a group of breeding birds from Louisiana, eastern Texas, and Arkansas, with wintering birds from the Mexican states of Yucatán and Quintana Roo. This “central” population appears to represent the easternmost end of a genetic cline across the Interior breeding range, with samples from eastern Texas and western Arkansas falling in intermediate locations in PC space and returning relatively high levels of admixture in Structure analyses. Mean *F*_*st*_ in 3-population structure models was 0.11. Neither clustering method found geographically coherent clusters beyond *k*=3 (Supplementary Figure 3-4).

**Figure 1:**
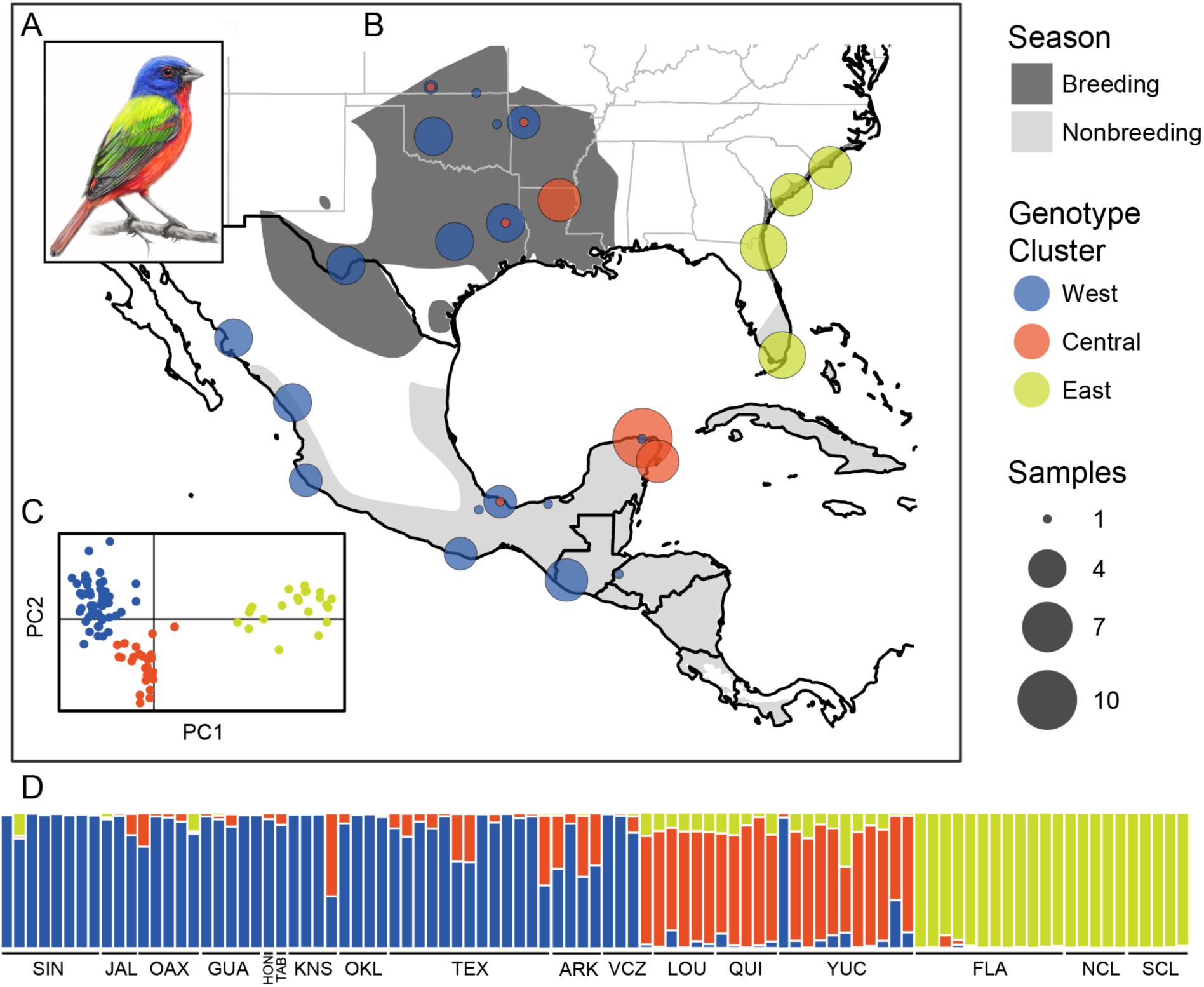
(A) Male *P. ciris*. (B) Sampling localities with points colored by *k*-means cluster and scaled to the number of samples. (C) Sample coordinates and k-means clusters on the first two PC axes. (D) Structure results at *k*=3, with each vertical bar representing a sample and the colors depicting the proportion of inferred ancestry from each population. Locality abbreviations are: SIN (Sinaloa), JAL (Jalisco), OAX (Oaxaca), GUA (Guatemala), HON (Honduras), TAB (Tabasco), KNS (Kansas), OKL (Oklahoma), TEX (Texas), ARK (Arkansas), VCZ (Veracruz), LOU (Louisiana), QUI (Quintana Roo), YUC (Yucatán), FLA (Florida), NCL (North Carolina), SCL (South Carolina).

Discriminant analyses estimated assignment probabilities over 0.99 for all individuals in both 2- and 3-population models. In cross-validation analyses, DAPC models trained on a random sample of half the individuals in each population correctly predicted the population assignment of an average of 99.7% of the remaining individuals at *k=*2, and 97.4% at *k*=3. DAPC cross-validation was also surprisingly robust at *k* values of 4 (85.5%) and 5 (76.2%); suggesting that denser population sampling could reveal further genetic structure in the interior breeding range.

### MtDNA

The population assignment scheme that explained the highest percentage of total variance in AMOVA’s grouped samples from Cuba and the Bahamas with eastern populations, and those from Costa Rica and Nicaragua with western populations (Table 1). Models including a central population of Louisiana and Yucatán birds were consistently lower ranked than two-population models, but the highest-ranked 3-population model followed the same assignment scheme as the top model overall. All AMOVA’s were highly significant (*p*<0.01). Haplotype networks were similar to those inferred using the breeding-season dataset of Herr et al. 2006, with the majority of western samples sharing a single common haplotype, and most eastern samples sharing one of two alternate haplotypes (Supplementary Figure 5).

**Table 1:** Individual assignment schemes for mitochondrial DNA AMOVA’s, ranked by the percentage of total variance explained by the population factor. Column *K* gives the total number of populations in the model. Columns with place names list a letter indicating the population assignment for each model.

| % Variation Explained by Population | *k* | Cuba | Bahamas | Central America |
| --- | --- | --- | --- | --- |
| 32.38 | 2 | E | E | W |
| 30.30 | 2 | W | E | W |
| 30.10 | 2 | E | W | W |
| 28.57 | 3 | E | E | W |
| 28.41 | 2 | W | W | W |
| 28.37 | 2 | E | E | E |
| 26.82 | 3 | C | E | W |
| 25.87 | 2 | W | E | E |
| 25.67 | 2 | E | W | E |
| 25.31 | 3 | E | E | E |
| 23.53 | 2 | W | W | E |
| 23.45 | 3 | C | E | E |

### Demographic Modeling

The west-central model had the lowest AIC score, but the difference in AIC scores between models was only 0.78; suggesting that the west-central and east-central models were equally supported given the data (Burnham and Anderson 2002). In both models the internode distance between the first and second divergence events is relatively short (around 17% of the total tree depth), and migration is much higher between west and central than east and central populations after the last divergence (Table 2, Supplementary Figure 6-7).

**Table 2:** δaδi parameter estimates and 95% confidence intervals for 3-population IM models with west+central or east+central populations as sister taxa. Migration parameters (“Nm_”) are the number of migrants per generation, with the receiving population listed first. “Ll_model” is the log-likelihood of the model under optimized parameter sets.

|  | West - Central | East - Central |
| --- | --- | --- |
| **N<sub>ref</sub>** | 92,806 (81,759 – 104,907) | 116,157 (102,160 – 122,264) |
| **N<sub>w</sub>** | 592,365 (529,869 – 695,108) | 572,601 (520,918 – 678,487) |
| **N<sub>e</sub>** | 43,702 (33,913 – 56,992) | 41,627 (30,303 – 50,487) |
| **N<sub>c</sub>** | 256,213 (204,475 – 322,629) | 304,896 (261,781 – 370,534) |
| **T1** | 566,477 (524,704 – 606,666) | 461,452 (418,442 – 537,264) |
| **T2** | 646,705 (605,602 – 684,570) | 680,303 (611,003 – 742,773) |
| **Nm<sub>wc</sub>** | 7.12 (5.51 – 8.81) | 6.95 (5.61 – 9.14) |
| **Nm<sub>cw</sub>** | 3.75 (3.08 – 4.58) | 3.57 (2.69 – 4.08) |
| **Nm<sub>ec</sub>** | 0.93 (0.87 – 1.05) | 0.98 (0.90 – 1.06) |
| **Nm<sub>ce</sub>** | 1.82 (1.43 – 2.7) | 2.00 (1.41 – 2.39) |
| **ll<sub>model</sub>** | -620.58 (-660.74 – -607.95) | -620.97 (-659.70 – -612.73) |

In the highest-likelihood parameter set, eastern and west+central populations diverge approximately 646,000 years ago, followed by central and west populations approximately 566,000 years ago. Effective population sizes scale with census sizes derived from BBS data (Blancher et al., 2007), with an estimated Ne/Nc for all populations combined of 0.081. Gene flow is highest between Central and West populations (3-9 migrants/generation), but also significant between East and Central (1-3 migrants/generation). Although we observed relatively low uncertainty across bootstrap replicates, exact figures for divergence times and population sizes should be interpreted with some caution given uncertainty in the generation time and genome-wide mutation rate.

### Morphology

As in Storer 1951 and Thompson 1991, we observed a cline in wing length across Texas, Oklahoma, Arkansas, and Louisiana (Figure 2; Supplementary Table 3), with the shortest-winged specimens collected in Louisiana. In regression analyses, wing chord (*p*<0.01, *R*^2^=0.33) and the first PC axis of wing chord and tarsus length (*p*<0.01, *R*^2^=0.29) were significantly correlated with longitude. The regression slope for tarsus length alone was not significant (*p=*0.07, *R*^2^=0.04). For the 21 specimens with both genomic and morphological data, wing chord (*p*<0.01, *R*^2^=0.38) and wing/tarsus PC1 (*p*<0.01, *R*^2^=0.24), but not tarsus length (*p*=0.67, *R*^2^=0.01), were also significantly correlated with the proportion of “central” ancestry in Structure results.

**Figure 2:**
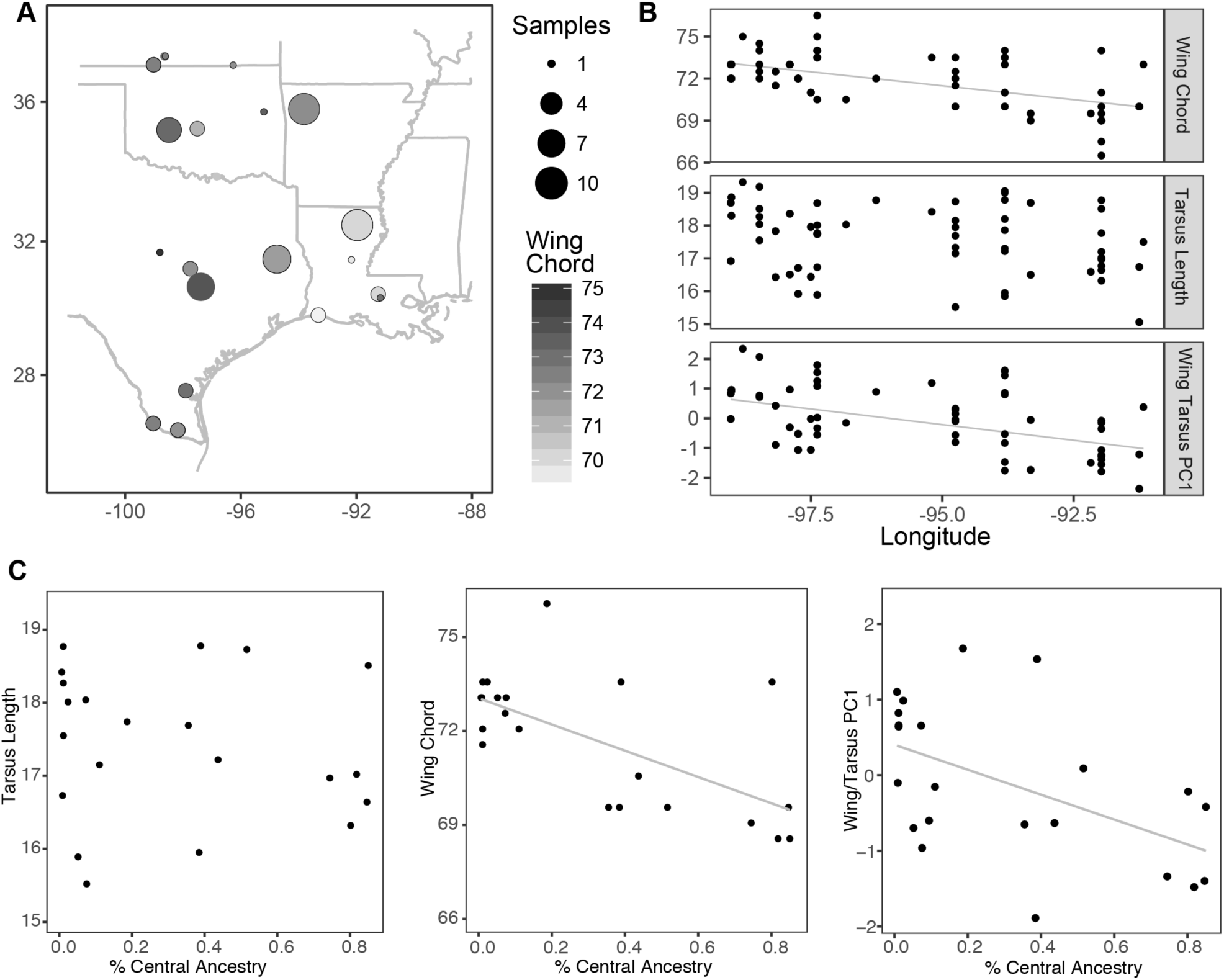
(A) map of specimen localities with points scaled by sample size and color graded by mean wing chord. (B) Linear regression of morphology as a function of longitude, with significant correlations shown as grey lines. (C) Linear regression of morphology as a function of the proportion of “Central” ancestry in 3-population Structure runs.

## Discussion

### Migratory Connectivity

Our study used thousands of genome-wide SNPs along with mtDNA sequences to produce the first range-wide map of migratory connectivity in Painted Buntings (Figure 1), yielding two major findings. First, we show that the Atlantic and interior breeding populations maintain allopatry year-round. Our clustering analyses failed to group any individuals from the Atlantic Coast breeding population with any individuals from a wintering localities in Mexico or Central America, a result consistent with previously hypothesized wintering range limits (Thompson, 1991). Second, we found strong signals of a migratory divide that corresponds with the geographic break between *P. c. pallidior* and *P. c. ciris* in eastern Texas proposed by Storer (1951). These genetically-conserved migratory programs may have important implications for the role of seasonal migration in shaping the evolutionary trajectory of populations. While seasonal migration may alternately facilitate gene flow and promote homogenization, or restrict gene flow and promote differentiation (Arguedas and Parker, 2000; Baker et al., 1994; Rohwer and Irwin, 2011), our data support the latter hypothesis in Painted Buntings, potentially indicating the presence incipient species. We believe this pattern reflects the consequences of extreme site fidelity across both the breeding and wintering range of the species.

Even in relatively well-studied taxa such as birds, the difficulty of tracking individuals and populations year-round has impeded research on many aspects of the ecology and evolution of migratory behavior (Webster et al., 2002). Painted Buntings are no exception, with previous studies proposing alternate migratory routes but failing to provide conclusive evidence of range-wide patterns. Based on similarities in plumage brightness and wing length, Storer (1951, 1982) concluded that individuals breeding in the Mississippi Valley migrated directly across the Gulf of Mexico to Winter on the Yucatan Peninsula. Prior to the present study, however, only a small number of mist-net captures on a single barrier island (Simons et al., 2004) and anecdotal observations from ships on trans-Gulf crossings (e.g. Frazar 1881) or from oil platforms (Sullivan et al., 2009) provided support for this trans-Gulf migratory route. Similarly, using banding records and differences in mean wing length between populations, Thompson (1991) proposed eastern Painted Buntings winter exclusively in southern Florida and the Caribbean, with western Painted Buntings wintering across Mexico and Central America. Unfortunately, a geolocator study (Contina et al., 2013) attempting in part to verify Thompson’s hypotheses was hindered by low retrieval rates and stochastic individual behavior. By resolving these longstanding questions in Painted Bunting biology, our work joins recent studies by Delmore et al. (2014, 2016) and others in highlighting the utility of genetic data to address recalcitrant problems in the natural history of avian migration.

### Phylogeography

The discordance between the longitude of the Painted Bunting subspecies boundary (recognized on the basis of morphology), and the longitudinal limits of allopatric Atlantic and interior breeding populations has long vexed ornithologists (e.g. Thompson 1991, Herr et al. 2011). Are current subspecies range limits an accurate reflection of phylogeographic structure and demographic independence, or do plumage and wing length characteristics reveal a hidden history of assortative mating and extinction? Our research corroborates both hypotheses. While our results broadly corroborate the sole previous study of genetic variation within *Passerina ciris* based on a mtDNA marker (Herr et al. 2011), the increased resolution afforded by genome-wide SNP data also reveals a previously undiagnosed genetic cluster consistent with morphological work by Storer (1951) and Thompson (1991) (Figure 1). We note that while our two methods of clustering analysis –*k-*means clustering and the Bayesian model-based method Structure -- produced conflicting results for the optimal number of populations in our sampling, individual membership assignments were concordant between programs when holding the value of K constant.

We suggest that the contemporary genetic structure of the Painted Bunting is likely the result of one of three scenarios. Based on clinal variation in wing length across the interior population and niche modeling that suggests the species’ east / west range gap is well within its potential climatic niche (Shipley et al. 2013), adaptive morphological evolution related to migratory distance may have been followed by the extinction of an intermediate portion of the ancestral range, potentially due to the fitness costs of trans-Gulf migration. Alternatively, the gap may be the result of disruption followed by slow interglacial population expansion from refugia during the Pleistocene (Johnson et al., 2004), or a stepping stone colonization event followed by preservation of the primary migratory axis (Greenberg and Marra 2005).

### Implications for Conservation

Painted Buntings are a charismatic species that, at least along the Atlantic coast, is restricted to a narrow strip of habitat heavily impacted by agricultural and residential development, and have attracted substantial conservation efforts. The species is listed as “Near Threatened” on the IUCN red list, and is considered a “species of special concern” in the US Fish and Wildlife Service’s Migratory Bird Program Strategic Plan (USFWS 2008). These designations are based primarily on data from the Breeding Bird Survey (BBS) finding that eastern populations have declined at an average rate of −1.17%/yr (95% CI −3.12, 0.08; Sauer et al., 2015), though trends in the region are not well supported because relatively few BBS routes are located in suitable habitat. In the west, BBS trends are highly variable, with populations apparently stable or expanding across northern Texas, Arkansas, and Oklahoma, while declining in Louisiana and Mississippi.

In contrast to earlier hypotheses of population structure in the species, which split eastern and western subspecies either in eastern Texas (Storer 1951) or across the breeding range gap in the Southeastern US (Thompson 1991, Herr et al. 20011), our analysis identified three populations (Figure 1). We estimate that Eastern, Central, and Western populations have persisted as separate lineages for at least 500,000 years; exchanging migrants at rates high enough to mitigate genetic divergence in an evolutionary context (up to 9 migrants/generation between Western and Central populations), but too low to impact demographic rates on timescales relevant for conservation.

Re-examined in this framework, BBS survey data suggest that western populations are healthy and expanding in the north while central and eastern populations are declining. In Louisiana, BBS data identifies a well-supported decline of −1.85/yr (95% CI −2.95, −0.95) – a faster rate than in the east, where most conservation attention has focused. Eastern populations, meanwhile, have both the lowest effective population size and the lowest levels of gene flow with other populations. Though we do not agree with Thompson’s 1991 hypothesis that they represent a separate species, the eastern population would easily qualify as a distinct population segment in the context of Endangered Species Act listing criteria if abundance declined significantly in the future. However, populations in the Mississippi alluvial valley region are also genetically differentiated from other Painted Buntings and show a more well-supported trend of population declines than eastern birds (mean −1.70/yr, 95%CI −2.86, −0.59), and should be monitored as a distinct demographic unit when analyzing survey data for the species.

## Conclusions

We documented a migratory divide in the Painted Bunting using mitochondrial DNA, genome-wide SNPs, and morphological analyses. Breeding populations from Louisiana largely migrate to the Yucatán Peninsula, while those from Central Texas and Oklahoma migrate first to molting grounds in western Mexico and then to southern Mexico and Central America. Genetic data found that the Atlantic coast breeding population is allopatric from interior populations year-round; wintering only in southern Florida, the Bahamas, and Cuba. These populations have a deep history of divergence with gene flow, with all three splitting approximately 500,000-700,000 years ago but continuing to exchange an average of 1-9 migrants per generation after divergence. The genetic cline west of the Mississippi is also associated with variation in wing length; suggesting that selection may be promoting morphological divergence in populations with different migration strategies.

It is remarkable that basic life history traits of this charismatic and relatively well-studied species remain to be discovered, and points to an ongoing need for natural history observations to drive advances in both conservation and evolutionary biology. In this case our results support monitoring of the putatively declining central and eastern populations as separate demographic and evolutionary units for conservation purposes. This species is also a promising system for further studies of the genetic mechanisms underlying variation in molt and migration in songbirds. Painted Buntings were historically a common caged pet in the United States and have reportedly been bred in captivity (Greene, 1883), making them a potentially tractable system for conducting controlled crosses. Because Atlantic Coast and Interior breeding populations differ in the timing and location of molt in addition to migration (before vs. during fall migration, respectively), future studies in this system could provide insight into the mechanisms underlying temporal variation in the full annual cycle of Passerine birds.

## Acknowledgements

We thank the staff and curators of the Burke Museum of Natural History and Universidad Nacional Autónoma de México for assistance with tissue loans, without which this work would not have been possible. <additional acknowledgements pending review>

## Data Availability

Complete data to reproduce the analyses described in this manuscript can be found at: (dryad link pending submission; temporary dropbox link: https://www.dropbox.com/sh/r4i55p09ea47uyt/AAA-DOzXhDAbxxVQlm5NpJffa?dl=0)

---

**Supplementary Figure 1:**
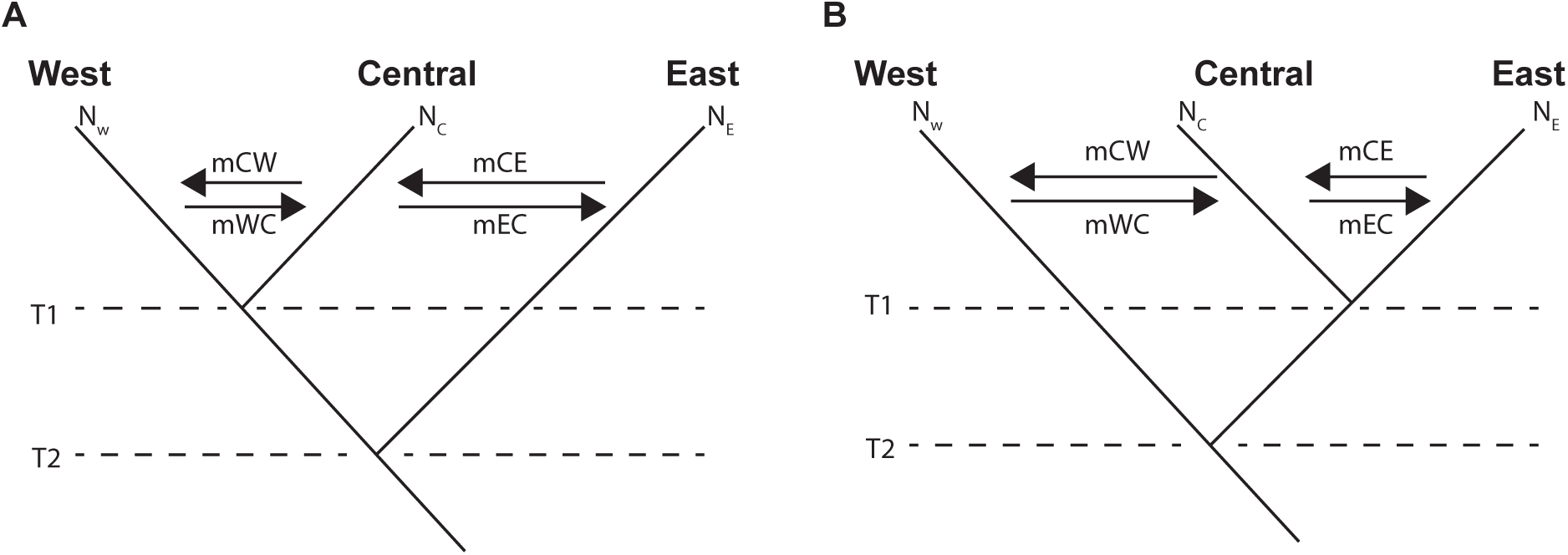
Demographic models compared in δaδi, with West and Central populations (A) or East and Central populations (B) as sister taxa.

**Supplementary Figure 2:**
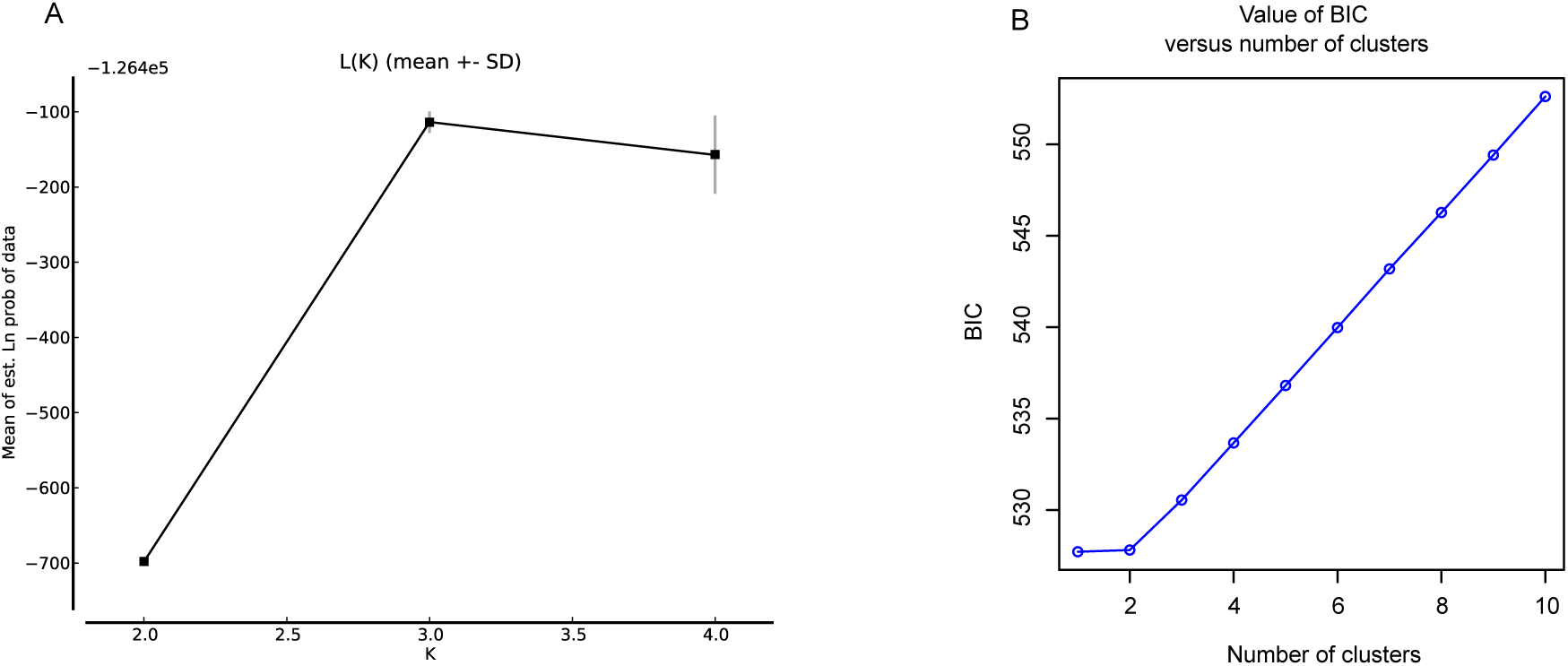
(A) Structure mean marginal likelihood and (B) *k-means* BIC for *k*=2-4.

**Supplementary Figure 3:**
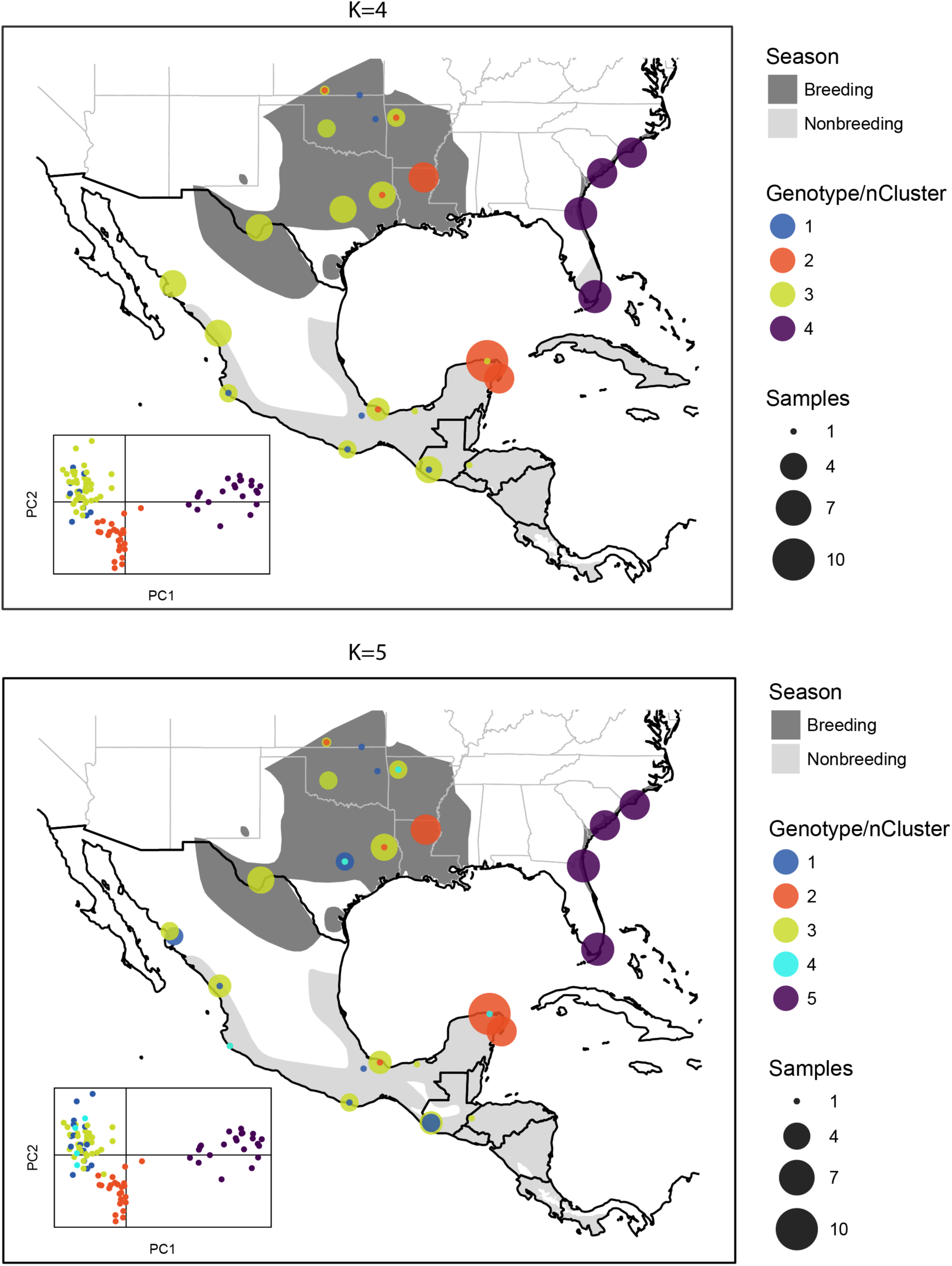
Map and PCA plot of *k*-means clustering on genotype data for *k*=4-5.

**Supplementary Figure 4:**
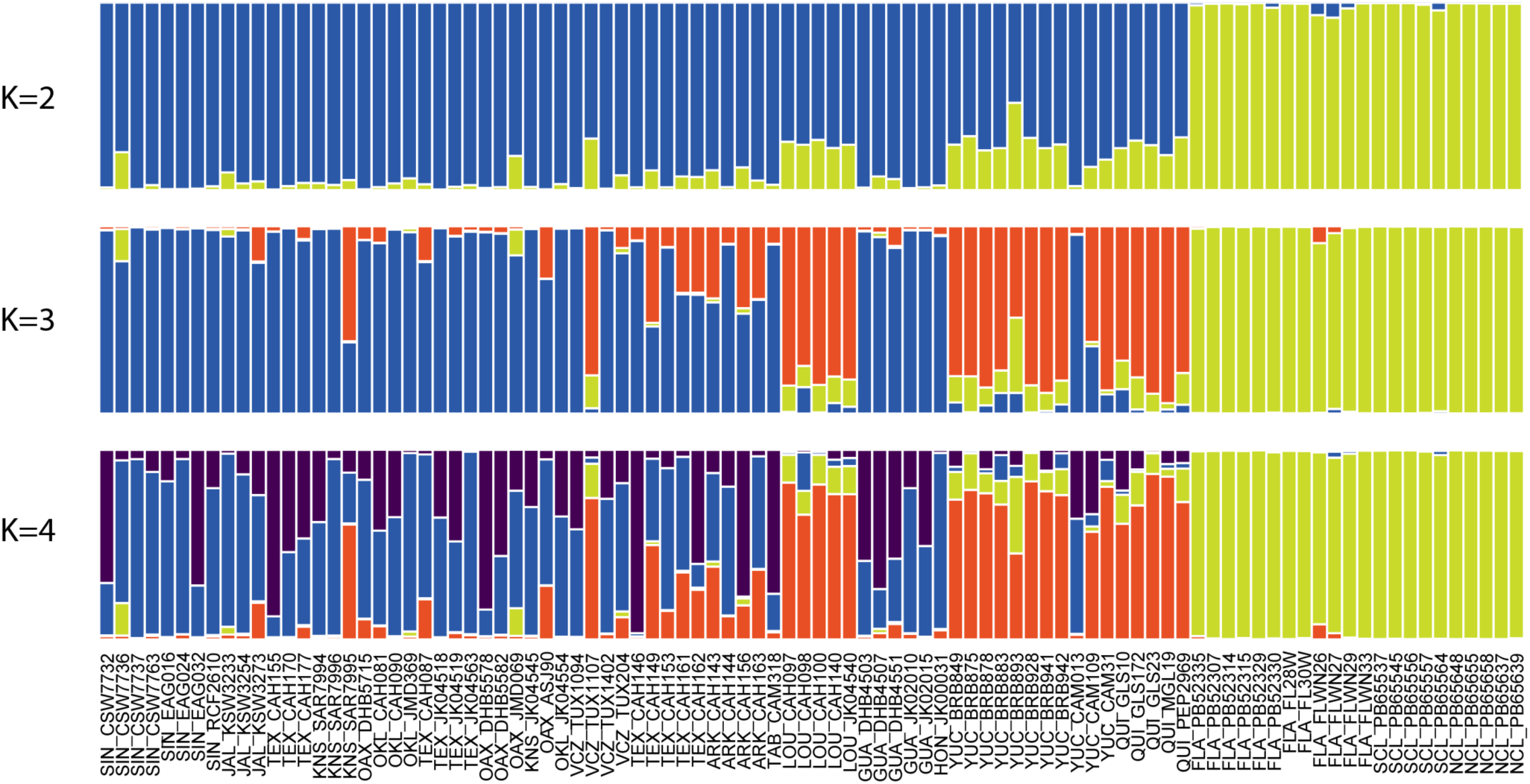
Average Structure results across five runs for each value of *k* (via CLUMPP), for *k*=2-4. Samples are ordered horizontally by longitude.

**Supplementary Figure 5:**
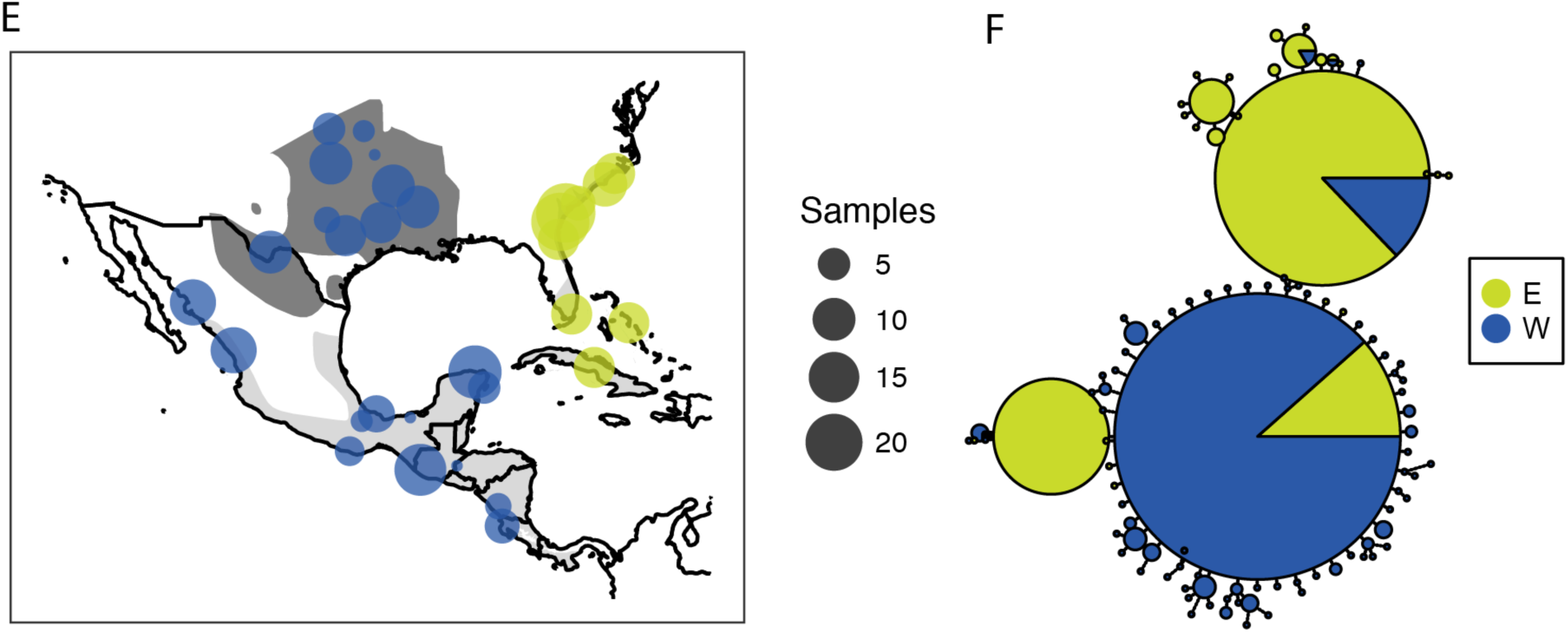
Mitochondrial DNA results. Map of sampling localities (E) and median-joining haplotype network (F), colored by the best-performing AMOVA assignment model.

**Supplementary Figure 6:**
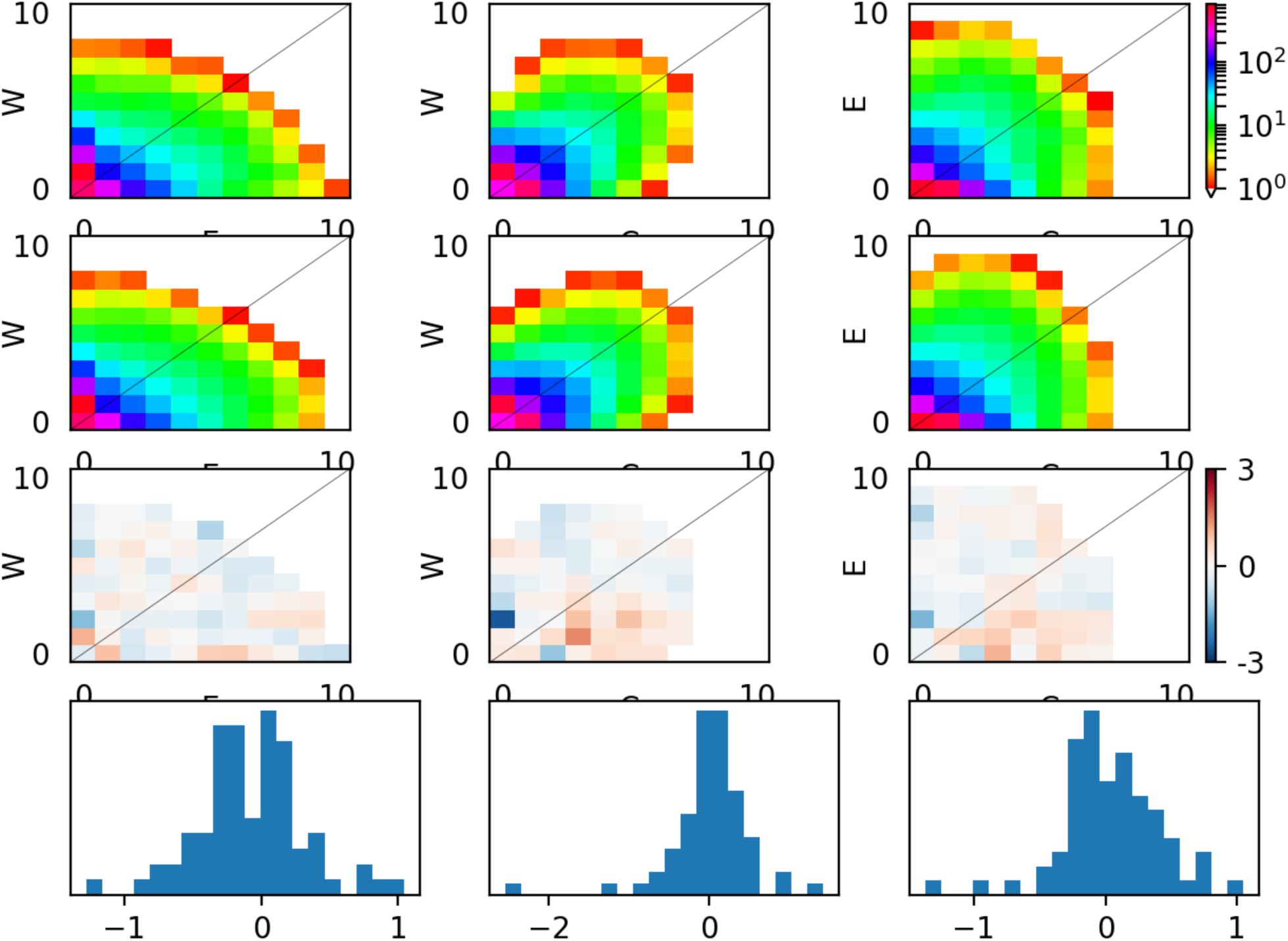
δaδi observed SFS (first row), model SFS (second row) and residuals (third row) for the western+central 3-population IM model.

**Supplementary Figure 7:**
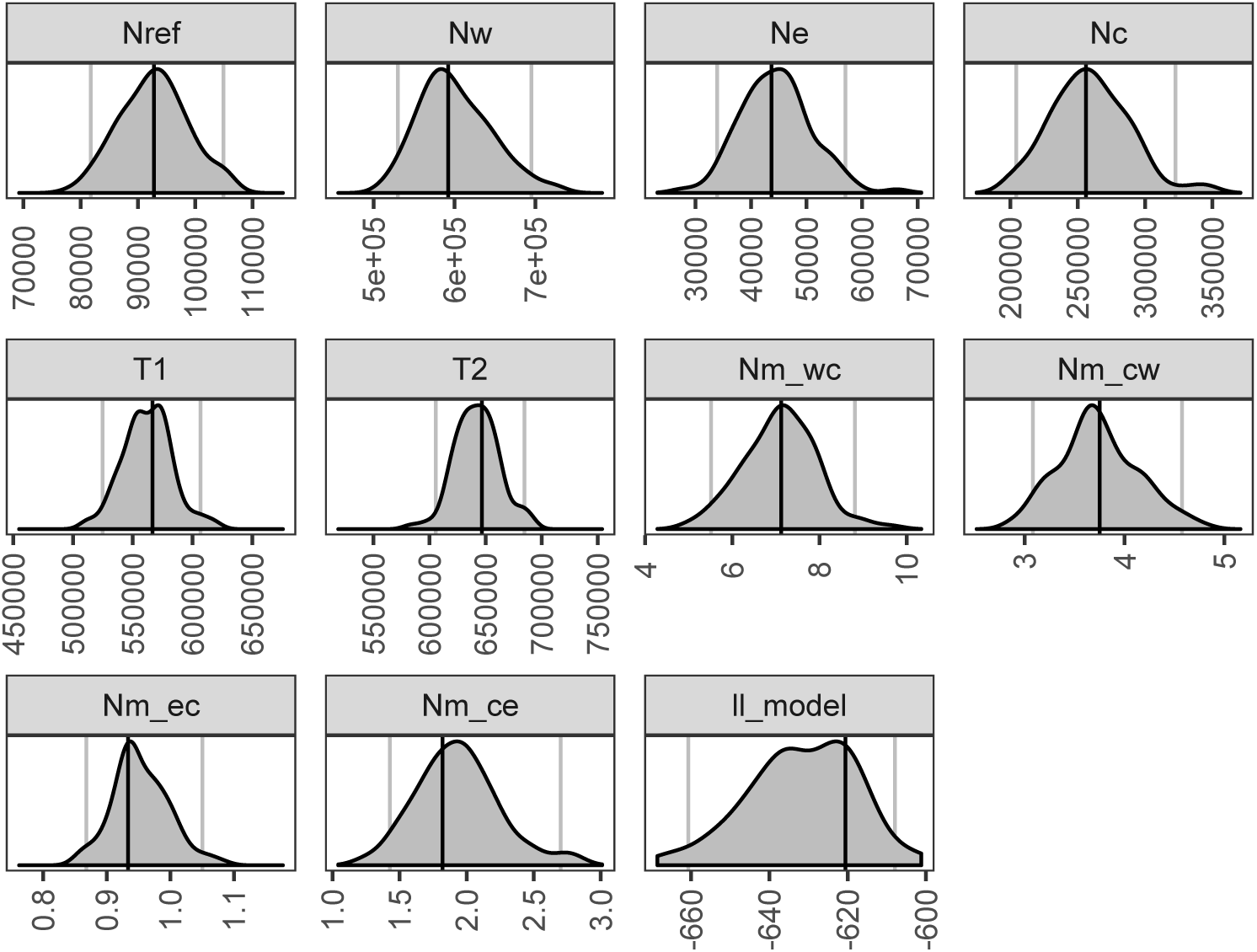
δaδi demographic parameter estimates. Distributions give probability densities across 100 parametric bootstrap replicates. Vertical lines show 95% confidence intervals (grey) and optimal (black) parameter values.

**Supplementary Table 1:** Specimen and sample information for ddRADseq analyses. Bolded samples were used in demographic modeling.

| Sample ID | State | Latitude | Longitude | Source | Museum Number | MO | DY | YR |
| --- | --- | --- | --- | --- | --- | --- | --- | --- |
| **ARK_CAH143** | **Arkansas** | **35.80** | **-93.81** | **UWBM** | **101739** | **6** | **15** | **2005** |
| **ARK_CAH144** | **Arkansas** | **35.80** | **-93.81** | **UWBM** | **101740** | **6** | **15** | **2005** |
| ARK_CAH156 | Arkansas | 35.80 | -93.81 | UWBM | 101752 | 6 | 14 | 2005 |
| **ARK_CAH163** | **Arkansas** | **35.80** | **-93.81** | **UWBM** | **101758** | **6** | **14** | **2005** |
| FLA_FL28W | Florida | 25.49 | -80.47 | USGS | FL28W | 1 | 23 | 2004 |
| **FLA_FL30W** | **Florida** | **25.49** | **-80.47** | **USGS** | **FL30W** | **1** | **23** | **2004** |
| FLA_FLWN26 | Florida | 25.49 | -80.47 | USGS | FL_W_N26 | 1 | 23 | 2004 |
| FLA_FLWN27 | Florida | 25.49 | -80.47 | USGS | FL_W_N27 | 1 | 23 | 2004 |
| **FLA_FLWN29** | **Florida** | **25.49** | **-80.47** | **USGS** | **FL_W_N29** | **1** | **23** | **2004** |
| FLA_FLWN33 | Florida | 25.49 | -80.47 | USGS | FL_W_N33 | 1 | 23 | 2004 |
| **FLA_PB52307** | **Florida** | **30.44** | **-81.44** | **USGS** | **PB52307** | **8** | **2** | **2000** |
| FLA_PB52314 | Florida | 30.44 | -81.44 | USGS | PB52314 | 8 | 2 | 2000 |
| **FLA_PB52315** | **Florida** | **30.44** | **-81.44** | **USGS** | **PB52315** | **8** | **2** | **2000** |
| FLA_PB52329 | Florida | 30.41 | -81.43 | USGS | PB52329 | 8 | 3 | 2000 |
| FLA_PB52330 | Florida | 30.41 | -81.43 | USGS | PB52330 | 8 | 3 | 2000 |
| **FLA_PB52335** | **Florida** | **30.44** | **-81.47** | **USGS** | **PB52335** | **8** | **4** | **2000** |
| **GUA_DHB4503** | **Retalhuleu** | **14.60** | **-91.61** | **UWBM** | **104172** | **1** | **10** | **2002** |
| GUA_DHB4507 | Retalhuleu | 14.60 | -91.61 | UWBM | 104176 | 1 | 10 | 2002 |
| GUA_DHB4551 | Retalhuleu | 14.60 | -91.61 | UWBM | 104220 | 1 | 10 | 2002 |
| **GUA_JK02010** | **Retalhuleu** | **14.60** | **-91.61** | **UWBM** | **109863** | **1** | **10** | **2002** |
| GUA_JK02015 | Retalhuleu | 14.60 | -91.61 | UWBM | 109868 | 1 | 11 | 2002 |
| HON_JK00031 | Copán | 14.90 | -88.90 | UWBM | 94126 | 4 | 15 | 2000 |
| JAL_KSW3233 | Jalisco | 19.52 | -105.07 | UNAM | KSW 3233 | 10 | 18 | 1999 |
| JAL_KSW3254 | Jalisco | 19.52 | -105.07 | UNAM | KSW 3254 | 10 | 21 | 1999 |
| **JAL_KSW3273** | **Jalisco** | **19.52** | **-105.07** | **UNAM** | **KSW 3273** | **10** | **22** | **1999** |
| KNS_JK04545 | Kansas | 37.02 | -96.26 | UWBM | 111618 | 5 | 28 | 2004 |
| KNS_SAR7994 | Kansas | 37.26 | -98.65 | UWBM | 90011 | 6 | 19 | 2009 |
| KNS_SAR7995 | Kansas | 37.27 | -98.61 | UWBM | 90012 | 6 | 19 | 2009 |
| KNS_SAR7996 | Kansas | 37.26 | -98.62 | UWBM | 90013 | 6 | 21 | 2009 |
| **LOU_CAH097** | **Louisiana** | **32.48** | **-91.97** | **UWBM** | **101694** | **6** | **2** | **2004** |
| LOU_CAH098 | Louisiana | 32.48 | -91.97 | UWBM | 101695 | 6 | 2 | 2004 |
| LOU_CAH100 | Louisiana | 32.48 | -91.97 | UWBM | 101697 | 6 | 2 | 2004 |
| LOU_CAH140 | Louisiana | 32.48 | -91.97 | UWBM | 101736 | 6 | 9 | 2005 |
| LOU_JK04540 | Louisiana | 32.48 | -91.97 | UWBM | 111613 | 6 | 2 | 2004 |
| **NCL_PB65637** | **North Carolina** | **33.86** | **-77.99** | **USGS** | **PB65637** | **8** | **28** | **2000** |
| NCL_PB65639 | North Carolina | 33.86 | -77.99 | USGS | PB65639 | 8 | 28 | 2000 |
| NCL_PB65648 | North Carolina | 33.87 | -78.00 | USGS | PB65648 | 8 | 29 | 2000 |
| **NCL_PB65655** | **North Carolina** | **33.87** | **-78.00** | **USGS** | **PB65655** | **8** | **29** | **2000** |
| **NCL_PB65658** | **North Carolina** | **33.87** | **-78.00** | **USGS** | **PB65658** | **8** | **29** | **2000** |
| OAX_ASJ90 | Oaxaca | 18.08 | -96.13 | UNAM | ASJ 90 | 4 | 14 | 1995 |
| OAX_DHB5578 | Oaxaca | 15.92 | -97.08 | UWBM | 105252 | 1 | 16 | 2004 |
| OAX_DHB5582 | Oaxaca | 15.92 | -97.08 | UWBM | 105256 | 1 | 16 | 2004 |
| OAX_DHB5715 | Oklahoma | 35.21 | -98.48 | UWBM | 105390 | 5 | 25 | 2004 |
| OAX_JMD069 | Oaxaca | 15.92 | -97.08 | UWBM | 108268 | 1 | 16 | 2004 |
| **OKL_CAH081** | **Oklahoma** | **35.21** | **-98.48** | **UWBM** | **101678** | **5** | **25** | **2004** |
| OKL_CAH090 | Oklahoma | 35.21 | -98.48 | UWBM | 101686 | 5 | 25 | 2004 |
| OKL_JK04554 | Oklahoma | 35.72 | -95.20 | UWBM | 111627 | 5 | 27 | 2004 |
| OKL_JMD369 | Oklahoma | 35.21 | -98.48 | UWBM | 108580 | 5 | 25 | 2004 |
| **QUI_GLS10** | **Quintana Roo** | **20.44** | **-86.90** | **UNAM** | **UNAM** | **1** | **13** | **1995** |
| QUI_GLS172 | Quintana Roo | 20.44 | -86.90 | UNAM | UNAM | 3 | 17 | 1995 |
| **QUI_GLS23** | **Quintana Roo** | **20.44** | **-86.90** | **UNAM** | **UNAM** | **1** | **15** | **1995** |
| **QUI_MGL19** | **Quintana Roo** | **20.44** | **-86.90** | **UNAM** | **UNAM** | **1** | **15** | **1995** |
| **QUI_PEP2969** | **Quintana Roo** | **20.44** | **-86.90** | **UNAM** | **UNAM** | **10** | **12** | **1994** |
| **SCL_PB65537** | **South Carolina** | **32.72** | **-79.99** | **USGS** | **PB65537** | **8** | **18** | **2000** |
| SCL_PB65545 | South Carolina | 32.72 | -79.99 | USGS | PB65545 | 8 | 18 | 2000 |
| **SCL_PB65556** | **South Carolina** | **32.72** | **-79.99** | **USGS** | **PB65556** | **8** | **18** | **2000** |
| SCL_PB65557 | South Carolina | 32.72 | -79.99 | USGS | PB65557 | 8 | 18 | 2000 |
| SCL_PB65564 | South Carolina | 32.72 | -79.99 | USGS | PB65564 | 8 | 18 | 2000 |
| SIN_CSW7732 | Sinaloa | 26.28 | -108.80 | UWBM | 90608 | 9 | 28 | 2010 |
| SIN_CSW7736 | Sinaloa | 26.28 | -108.80 | UWBM | 90612 | 9 | 28 | 2010 |
| SIN_CSW7737 | Sinaloa | 26.28 | -108.80 | UWBM | 90613 | 9 | 28 | 2010 |
| SIN_CSW7763 | Sinaloa | 26.28 | -108.80 | UWBM | 90637 | 9 | 30 | 2010 |
| **SIN_EAG016** | **Sinaloa** | **23.25** | **-105.75** | **UWBM** | **81416** | **7** | **17** | **2005** |
| SIN_EAG024 | Sinaloa | 23.25 | -105.75 | UWBM | 81424 | 7 | 18 | 2005 |
| SIN_EAG032 | Sinaloa | 23.25 | -105.75 | UWBM | 81432 | 7 | 18 | 2005 |
| SIN_RCF2610 | Sinaloa | 23.25 | -105.75 | UWBM | 81134 | 7 | 17 | 2005 |
| TAB_CAM318 | Tabasco | 18.36 | -92.56 | UNAM | CAM 318 | 3 | 11 | 1996 |
| TEX_BTS05071 | Texas | 29.59 | -103.00 | UWBM | 100190 | 5 | 28 | 2005 |
| TEX_CAH087 | Texas | 30.66 | -97.38 | UWBM | 101683 | 6 | 26 | 2004 |
| TEX_CAH146 | Texas | 31.48 | -94.75 | UWBM | 101742 | 6 | 7 | 2005 |
| TEX_CAH149 | Texas | 31.48 | -94.75 | UWBM | 101745 | 6 | 6 | 2005 |
| TEX_CAH153 | Texas | 31.48 | -94.75 | UWBM | 101749 | 6 | 6 | 2005 |
| TEX_CAH155 | Texas | 29.59 | -103.00 | UWBM | 101751 | 5 | 29 | 2005 |
| TEX_CAH161 | Texas | 31.48 | -94.75 | UWBM | 101756 | 6 | 6 | 2005 |
| TEX_CAH162 | Texas | 31.48 | -94.75 | UWBM | 101757 | 6 | 6 | 2005 |
| TEX_CAH170 | Texas | 29.59 | -103.00 | UWBM | 101765 | 5 | 29 | 2005 |
| **TEX_CAH177** | **Texas** | **29.59** | **-103.00** | **UWBM** | **101772** | **5** | **28** | **2005** |
| **TEX_JK04518** | **Texas** | **30.66** | **-97.38** | **UWBM** | **111592** | **6** | **26** | **2004** |
| TEX_JK04519 | Texas | 30.66 | -97.38 | UWBM | 111593 | 6 | 26 | 2004 |
| TEX_JK04563 | Texas | 30.66 | -97.38 | UWBM | 111636 | 6 | 27 | 2004 |
| VCZ_TUX1094 | Veracruz | 18.48 | -95.03 | UNAM | TUX 1094 | 10 | 18 | 1994 |
| **VCZ_TUX1107** | **Veracruz** | **18.48** | **-95.03** | **UNAM** | **TUX 1107** | **10** | **25** | **1994** |
| VCZ_TUX1402 | Veracruz | 18.48 | -95.03 | UNAM | TUX 1402 | 4 | 27 | 2003 |
| VCZ_TUX204 | Veracruz | 18.48 | -95.03 | UNAM | TUX 204 | 4 | 9 | 1994 |
| YUC_BRB849 | Yucatán | 21.52 | -87.70 | UWBM | 99969 | 1 | 13 | 2009 |
| YUC_BRB875 | Yucatán | 21.52 | -87.70 | UWBM | 99995 | 1 | 13 | 2009 |
| **YUC_BRB878** | **Yucatán** | **21.52** | **-87.70** | **UWBM** | **99998** | **1** | **14** | **2009** |
| YUC_BRB883 | Yucatán | 21.52 | -87.70 | UWBM | 100003 | 1 | 14 | 2009 |
| YUC_BRB893 | Yucatán | 21.52 | -87.70 | UWBM | 100013 | 1 | 14 | 2009 |
| YUC_BRB928 | Yucatán | 21.52 | -87.70 | UWBM | 100045 | 1 | 14 | 2009 |
| YUC_BRB941 | Yucatán | 21.52 | -87.70 | UWBM | 100057 | 1 | 15 | 2009 |
| **YUC_BRB942** | **Yucatán** | **21.52** | **-87.70** | **UWBM** | **100058** | **1** | **15** | **2009** |
| YUC_CAM013 | Yucatán | 21.51 | -87.67 | UNAM | CAM 013 | 3 | 9 | 1995 |
| **YUC_CAM109** | **Yucatán** | **21.51** | **-87.67** | **UNAM** | **CAM 109** | **3** | **19** | **1995** |
| **YUC_CAM31** | **Yucatán** | **21.51** | **-87.67** | **UNAM** | **CAM 31** | **3** | **10** | **1995** |

<see attached file>

**Supplementary Table 2:** Specimen and sample information for mtDNA analyses.

<see attached file>

**Supplementary Table 3:** Morphological data for interior breeding populations.

